# scTFBridge: A Disentangled Deep Generative Model Informed by TF-Motif Binding for Gene Regulation Inference in Single-Cell Multi-Omics

**DOI:** 10.1101/2025.01.16.633293

**Authors:** Feng-ao Wang, Ruikun He, Junwei Liu, Yixue Li

## Abstract

The interplay between transcription factors (TFs) and regulatory elements (REs) drives gene transcription, forming gene regulatory networks (GRNs). Advances in single-cell technologies now enable simultaneous measurement of RNA expression and chromatin accessibility, offering unprecedented opportunities for GRN inference at single-cell resolution. However, heterogeneity across omics layers complicates regulatory feature extraction. We present scTFBridge, a multi-omics deep generative model for GRN inference. scTFBridge disentangles latent spaces into shared and specific components across omics layers. By integrating TF-motif binding knowledge, scTFBridge aligns shared embeddings with specific TF regulatory activities, enhancing biological interpretability. Using explainability methods, scTFBridge computes regulatory scores for REs and TFs, enabling robust GRN inference. It consistently outperformed baseline methods in both cis- and trans-regulation inference tasks. Our results demonstrate that scTFBridge can uncover cell-type-specific susceptibility genes and distinct regulatory programs, offering new insights into gene regulation mechanisms at single-cell resolution.

## Main

Transcription of genes is primarily regulated by TFs, proteins that bind to specific DNA sequences to trigger diverse gene regulatory functions^1^. These regulatory elements (REs) are typically located in accessible chromatin regions, such as promoters, enhancers, and silencers^2,3^. The binding of TFs to REs and their spatial chromatin interaction with target genes (TGs) ultimately determines specific gene transcription processes^4^. The complex interplay among TFs, REs, and TGs can be represented as interconnected GRNs, the foundational elements of gene regulation in single cells^5-7^. Understanding the regulatory mechanisms within these GRNs is pivotal for elucidating how cellular identities are established and maintained. Such insights have far-reaching implications, from guiding cell fate engineering to informing strategies for disease prevention and treatment^8-10^.

Recent advances in single-cell high-throughput sequencing techniques have enabled the exploration of gene regulatory processes at single-cell resolution^11-13^. For example, single-cell RNA sequencing (scRNA-seq) facilitates the inference of cell-type–specific trans-regulatory interactions through computational tools such as DeepSEM^14^, PIDC^15^, GENIE3^16^, SupirFactor^17^, and SCENIC^18^. Similarly, single-cell assay for transposase-accessible chromatin using sequencing (scATAC-seq) provides insights into chromatin accessibility at specific gene regions (Peaks)^19^, enabling the identification of TF-RE interactions related to gene regulation through methods like SQUID^20^, DeepTFni^21^, and DirectNet^22^. However, these distinct omics layers are typically measured in separate cells, complicating efforts to directly link RNA expression and chromatin accessibility within the same cell^23^. Although unpaired data integration methods such as scJoint^24^, scGLUE^25^, IReNA^26^, SOMatic^27^, and UnpairReg^28^ have been developed to align and integrate these unpaired datasets, they are primarily optimized for tasks like cell annotation and data integration. As a result, their capacity to accurately capture the complex regulatory relationships within GRNs at single-cell resolution remains limited^29^.

The emergence of single-cell multi-omics technologies, such as 10x Multiome^30^ and SHARE-seq^31^, has transformed the field by enabling simultaneous profiling of paired RNA-seq and ATAC-seq data from the same cell. This advancement facilitates more accurate GRN inference and a deeper understanding of gene regulation at the single-cell level. However, integrating these high-dimensional and heterogeneous multi-omics datasets remains a significant challenge^32,33^. To address that, several computational methods, including MultiVI^34^, LINGER^35^, scButterfly^36^, UnitedNet^37^, and SCENIC+^38^, have been developed. These methods primarily employ multimodal alignment strategies, encoding different omics layers into a shared latent space to create a unified representation^39-42^. In many cases, adversarial learning with discriminators is used to align the distinct omics layers effectively. Despite these advances, a critical challenge persists: the “modality gap”^43,44^. This gap stems from the intrinsic heterogeneity of different omics layers, each capturing distinct aspects of complex biological processes^45^. Consequently, each modality retains unique private information that can impede the integration of a shared multi-omics latent space and limit accurate data reconstruction from the shared embedding^46,47^. Overcoming this modality gap is crucial for advancing multi-omics data integration and enabling more precise GRN analyses.

In this study, we propose scTFBridge, an interpretable deep generative model for multi-omics integration and GRN inference at the single-cell level. By leveraging mutual information theory^48-50^, scTFBridge minimizes mutual information and disentangles latent variables of both RNA-seq and ATAC-seq data into modality-shared and modality-private subspaces. The modality-shared representations are further aligned through contrastive learning to construct a unified latent space. Incorporating biological priors of TF-motif bindings, scTFBridge constrains latent variables to represent specific TF regulatory activities, effectively acting as a “bridge” between RNA-seq and ATAC-seq modalities. Using deep learning explainability techniques, we derive regulatory scores linking REs and latent TF variables to TG expressions during the generation of TG expression. Our results demonstrate that scTFBridge effectively disentangles modality-shared and modality-private information, yielding biologically interpretable embeddings that capture cell-type-specific TF regulatory activities. Benchmarking against state-of-the-art methods on single-cell cis- and trans-regulation datasets, scTFBridge consistently outperforms baseline approaches in regulatory feature extraction across diverse cell groups. In summary, scTFBridge provides an efficient computational framework for exploring phenotype-specific gene regulatory mechanisms at single-cell resolution. It advances multi-omics data integration and enhances our understanding of gene regulation dynamics in complex cellular systems.

## Results

### The framework of scTFBridge

scTFBridge is a biologically interpretable deep generative model designed for integrating single-cell multi-omics data and inferring cell-specific GRNs (Fig. 1a). It leverages a hybrid variational autoencoder^51^ (VAE) architecture that decomposes latent variables into modality-shared and modality-private components for each omics layer (Fig. 1b). Through mutual information theory and contrastive learning regularizations, scTFBridge effectively disentangles shared and private representations while aligning the shared latent space to capture regulatory signals common across multi-omics datasets (Fig. 1c, Methods). To enhance biological interpretability, scTFBridge incorporates prior knowledge by constraining the weights of ATAC-seq shared data decoder, promoting biologically meaningful embeddings in the shared latent space (Fig. 1d). Further enrichment analysis revealed the differential biological function of private/shared sets in cell-specific manner (Fig. 1e). Using model interpretation techniques, scTFBridge enables the extraction of RE and TF regulatory scores for TGs, enabling the establishment of single-cell-specific GRNs (Fig. 1f, g).

**Fig. 1.**
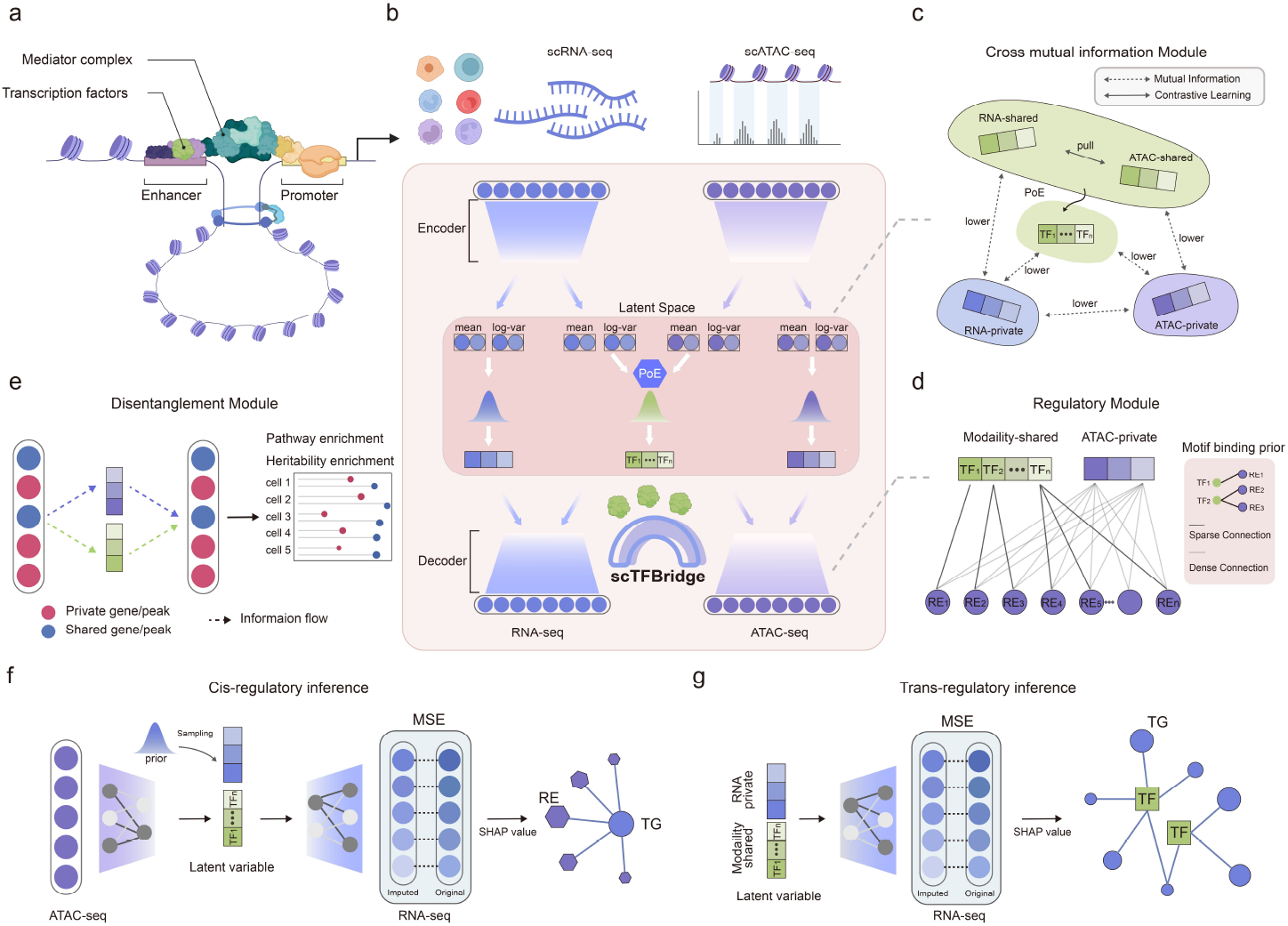
Overview of the scTFBridge framework. a. Schematic representation of RNA transcription, highlighting the cooperative roles of transcription factors (TFs) and regulatory elements (REs) in driving RNA synthesis. b. The model architecture of scTFBridge features a hybrid multi-modal variational autoencoder (VAE) designed for latent space dismantlement and representation of mutual information (MI) learning. c. Cross-mutual-information module for aligning modality-shared latent spaces and reducing MI between modality-shared and modality-private embeddings. d. Regulatory module in the ATAC decoder, where TF–motif binding constrains the connections between TF variables and regulatory elements. e. Definition of private/shared peak/gene sets according to their information flow from the input layer to disentangled latent space. Further enrichment analysis reveals the differential biological functions between private/shared sets in a cell-specific manner. f. Cis-regulation inference depicted for target gene (TG) expression prediction based on ATAC data, where the objective function minimizes the mean squared error (MSE) between original and reconstructed gene expression levels. g. Trans-regulation inference depicted for TG expression prediction using latent variables, employing the same computational strategy as in cis-regulation inference.

In detail, we developed a Cross Mutual Information (CMI) module to ensure a well-separated latent space across different modality domains (Fig. 1c). Independent neural networks were employed to estimate and minimize mutual information^52^ (MI) by introducing an upper bound^53^ on the MI between pairs of data domains (Methods). This regularization reduces MI not only between shared and private embeddings within each modality but also across RNA-private and ATAC-private embeddings, as well as between shared and modality-private embeddings (Fig. 1c). Additionally, a contrastive learning loss was incorporated to align shared embeddings of across modalities. This approach ensures that the shared latent space emphasizes TF–RE binding interactions and other regulatory signals common to RNA-seq and ATAC-seq while excluding modality-specific biological noise.

To preserve TF-RE regulatory relationships, we designed a regulatory module in scTFBridge that incorporates known TF–motif binding affinities within single-cell multi-omics data. In the shared embedding space, latent variables correspond to specific TFs. Prior knowledge of TF– motif binding constraints connections between these shared latent TF representations and the reconstructed REs (Fig. 1d, Methods). This regularization mechanism ensures that gene regulatory information from both RNA-seq and ATAC-seq omics is properly aligned and biologically meaningful. This regularization enhances interpretability by enabling direct examination of associations between multi-omics features and corresponding TF representations in the shared latent space.

To further elucidate gene regulatory mechanisms, we utilized SHAP (Shapley Additive Explanations) to quantify the contribution of input REs or shared latent TF variables to the TG expression reconstruction^54^ (Fig. 1f, g). SHAP scores were computed based on the mean squared error (MSE) between imputed and observed gene expression levels under specific feature perturbations (Methods). The biologically constrained model framework enables these SHAP scores to represent regulatory scores for each RE and TF linked to TGs. For GRN inference, we derived both cis-regulation (RE–TG) and trans-regulation (TF–TG) interactions across various cell types using these scores (Methods).

### scTFBridge facilitates cell-type-specific disentangled embedding learning

We applied scTFBridge to a single-cell multi-omics dataset comprising paired scRNA-seq and scATAC-seq measurements of bone marrow mononuclear cells (BMMCs) from healthy donors across multiple batches^55^. The learned latent embeddings of RNA-private and ATAC-private networks enabled visualization of domain-specific cell representations, alongside shared cell embeddings integrating both modalities. Uniform manifold approximation and projection (UMAP) of these embeddings highlighted distinct cellular information within each domain. In the RNA-private embedding space, most cell types clustered closely, except for erythroblasts and normoblasts, suggesting gene expression regulation independent of ATAC-seq signals during erythrocyte maturation (Fig. 2a). In contrast, the ATAC-private and shared embeddings captured additional aspects of major cellular heterogeneity (Fig. S1). To evaluate the scTFBridge’s capacity to learn cross-modal interactions, we conducted a TG expression reconstruction experiment using the learned shared embeddings with ATAC-seq data. Compared to baseline methods, scTFBridge demonstrated superior TG expression reconstruction accuracy (Fig. 2b, Fig. S2), indicating that paired ATAC-seq data effectively captured crucial gene expression signals.

**Fig. 2.**
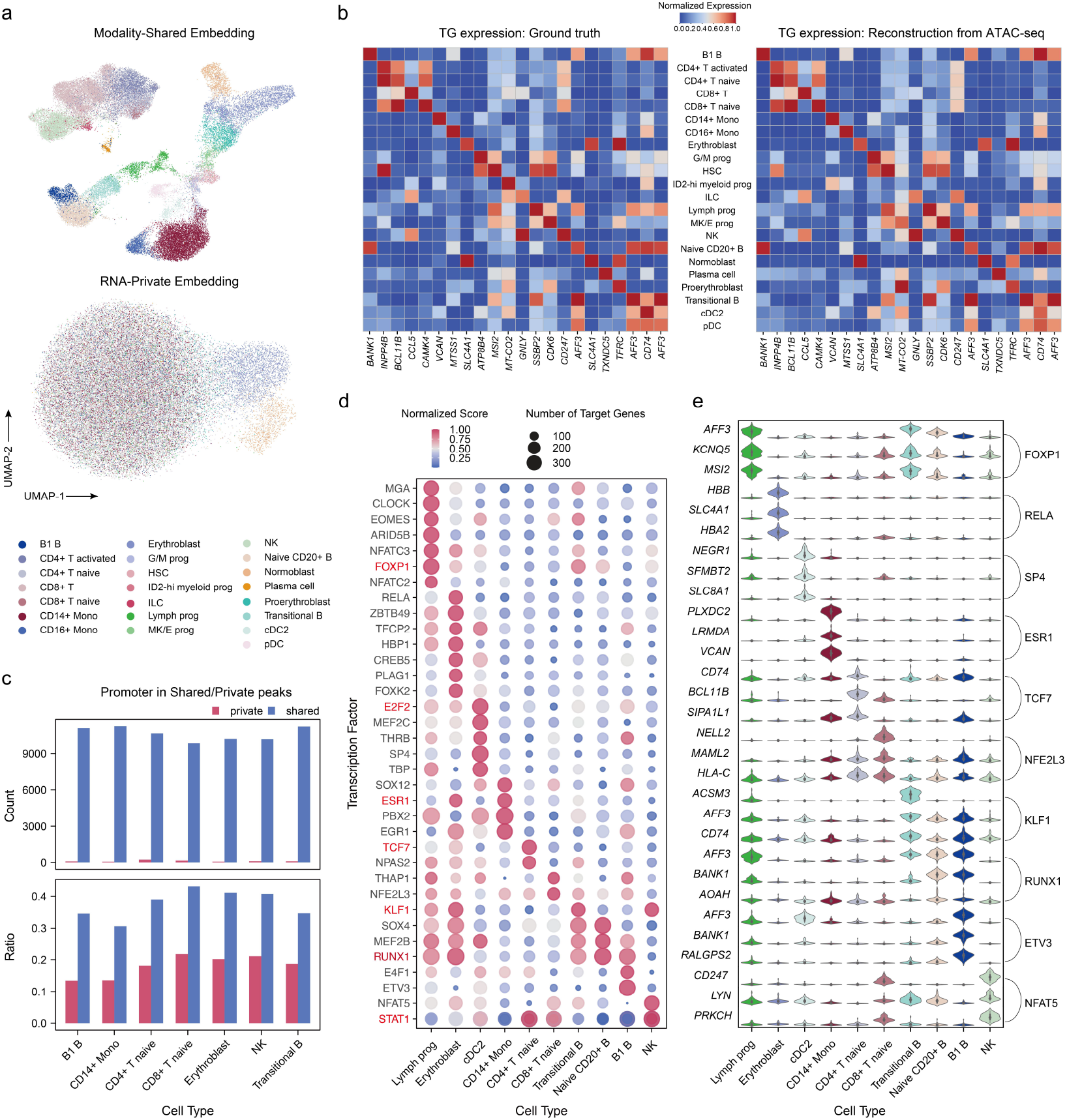
Interpretable TF-informed embedding learning of scTFBridge in BMMCs. a. UMAP visualization of modality-shared and RNA-private embeddings in bone marrow mononuclear cells (BMMCs). b. Comparison between ground-truth target gene (TG) RNA expression and scTFBridge-based reconstruction from ATAC-seq data. c. Quantification and ratio of annotated promoter peaks across shared and private peak sets for different cell types. d. Regulatory scores and target gene counts for cell-type-specific TF variables identified through biological interpretation analysis. e. Visualization of cell-type-specific gene regulons derived from high-ranked target genes (TGs) associated with TF variables.

We further assessed the contributions of different ATAC-seq data Peak features to the ATAC-shared and ATAC-private embeddings based on the information flow through different encoders. Peaks were categorized into shared or private groups by their relative contributions to shared or private embeddings (Methods). Comparing the number and ratio of promoter Peaks in each group, our result revealed that the modality-shared embedding primarily enriched gene regulatory signals (Fig. 2c). We then explored the cell-type-specific gene regulatory elements through biological interpretation analysis using scTFBridge (Methods). Our results identified cell-type specific TFs, such as ESR in cDC cells, TCF7 in CD4 naïve T cells, and SOX4 in B cells (Fig. 2d). Additionally, high-ranked TGs of TF variables revealed cell-type-specific gene regulons (Fig. 2e). These findings demonstrate scTFBridge’s capacity for biologically disentangled learning and its effectiveness in elucidating potential cellular gene regulatory mechanisms.

### scTFBridge reveals heritability enrichment of peaks in rheumatoid arthritis

We applied scTFBridge to a single-cell multi-omics dataset from profiled immune cells from rheumatoid arthritis tissues^56^. To visualize the latent representations, we used UMAP to project the modality-shared, RNA-private, and ATAC-private representations into a two-dimensional space (Fig. 3a). In the RNA-private embedding, immune cells formed close clusters, suggesting that most of the gene regulatory information for these cells in the RNA-seq data is captured by modality-shared embeddings, as compared to the ATAC-seq data. We then examined the mutual information (MI) between modality-shared and private representations (Fig. 3b). The MI between RNA-shared and ATAC-shared representations was higher than the MI between RNA-shared and RNA-private or ATAC-shared and RNA-private, demonstrating that scTFBridge effectively disentangles modality-private and modality-shared components.

**Fig. 3.**
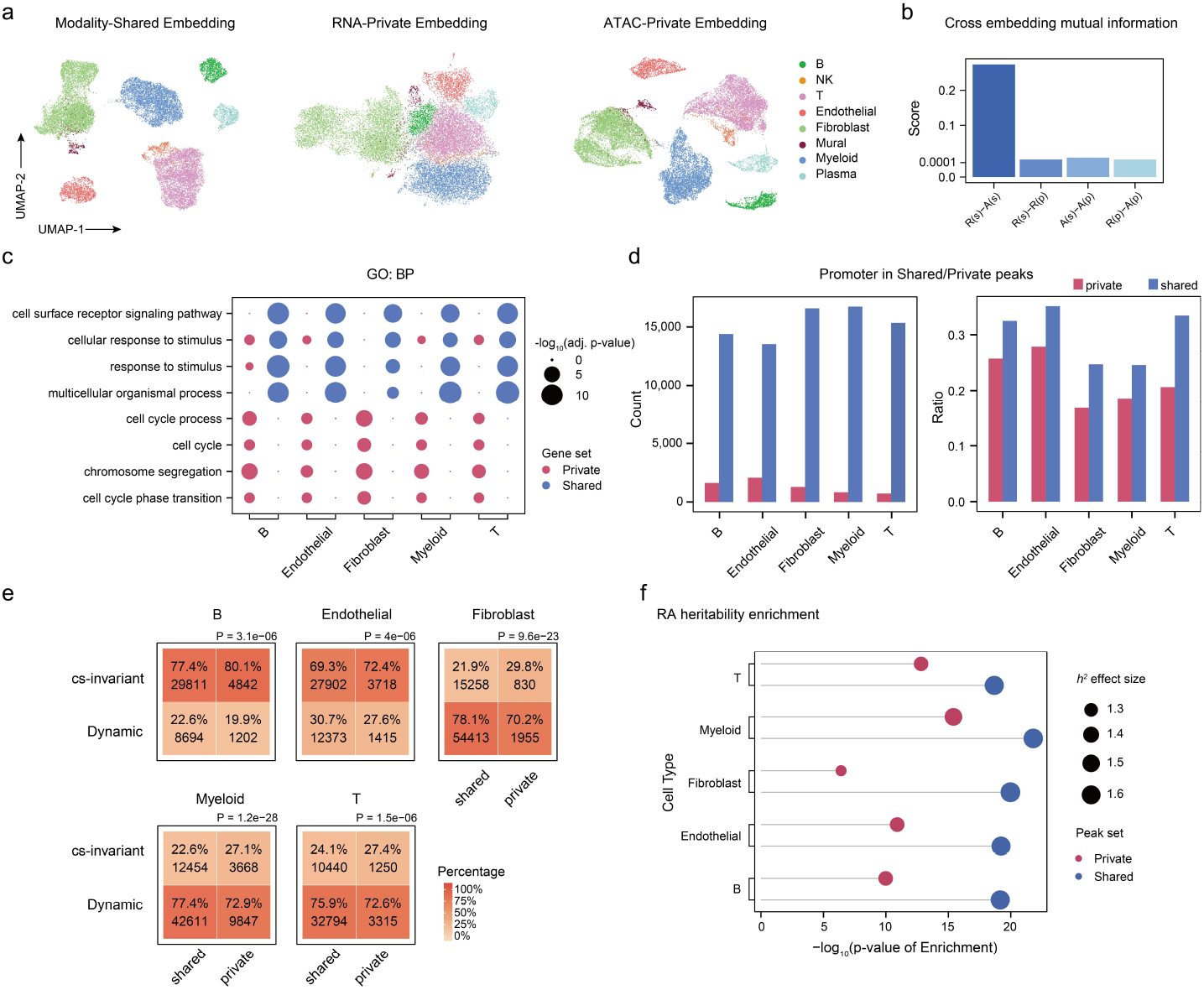
Identifying cell-specific modality-shared and private regulatory elements and functions in rheumatoid arthritis. a. UMAP visualization of modality-shared, RNA-private, and ATAC-private embedding in the dataset of rheumatoid arthritis. b. Mutual information (MI) comparison between different representations. “R” and “A” denote RNA and ATAC modalities, respectively; “s” and “p” denote shared and private embeddings, respectively. c. Gene Ontology (GO) pathway analysis comparing shared and private gene sets across different cell types. d. Quantification and ratio of annotated promoter peaks across shared and private peak sets in different cell types. e. Analysis of shared/private and dynamic/condition-specific (cs)-invariant peaks across cell types. Statistical significance was assessed using the Chi-square test, with p-values indicated. f. Heritability enrichment analysis of shared and private peak sets of single-cells from RA samples. The x-axis shows the -log_10_ enrichment p-value, and the dot size reflects the h^2^ effect.

To uncover deeper biological insights from distinct RNA-seq embeddings, we quantified the normalized absolute importance scores for each gene across different embeddings and identified the top 100 genes as their predominant gene programs (Methods). Gene Ontology (GO) pathway enrichment analysis revealed that the shared gene program primarily contributed to immune activation and receptor signaling, while the private gene program was associated with cell cycle regulation (Fig. 3c, Supplementary Data 1-10). Similarly, we assessed the enrichment of promoter peaks into modality-shared and ATAC-private embeddings. The results confirmed that the modality-shared embedding of scTFBridge captured a higher number of promoter peaks (Fig. 3d), highlighting its superior ability to capture gene regulatory signals in single-cell Multiome datasets.

To further evaluate the scTFBridge model’s ability to characterize crucial regulatory elements, we compared the shared- and ATAC-private predominated REs identified by scTFBridge with the dynamic and cell-state-invariant (cs-invariant) peaks identified in Gupta et al.^56^. Dynamic peaks were defined as those whose variation could be explained by paired RNA expression data, representing stronger gene regulatory effects^56^. In this comparison, we examined the overlap of peaks across two distinct groups (Fig. 3e). The results demonstrated that shared peaks were significantly enriched in the dynamic peak group (Chi-squared test, Methods). Additionally, we employed the stratified LD-score regression (S-LDSC)^57^ method to assess the heritability enrichment of shared and ATAC-private REs concerning GWAS-annotated summary statistics for rheumatoid arthritis (Methods). The enrichment scores revealed that shared peaks were more effective in capturing causal variants associated with the disease (Fig. 3f, Supplementary Data 11). These findings suggest that the scTFBridge model provides a more refined characterization of gene regulatory signals.

### Benchmarking analysis of cell-type-specific cis-regulatory inference

Accurate GRN inference remains challenging across different cell types and functional states due to the limited availability of gold-standard datasets and potential experimental biases. To assess the cell-type-specific cis-regulatory inference capabilities of scTFBridge, we integrated promoter-capture Hi-C (pcHi-C) data from three primary blood cell types^58^ and expression quantitative trait loci (eQTL) data from eight immune cell types^59,60^. These external validation datasets include SNPs significantly associated with gene expression, serving as known cis-regulatory RE-TG pairs for performance benchmarking (Supplementary Table 1, Methods). We compared scTFBridge’s performance in cis-regulatory inference against established methods, including PCC, Distance, Random, and SCENIC+^38^ (Methods).

Since the distance from transcription start sites (TSS) of TG is a critical factor in cis-regulatory inference, we categorized predicted RE-TG pairs into six distance groups, ranging from 0-5 kb to 100-200 kb, for evaluation. Using single-cell eQTL data as the ground truth, scTFBridge achieved the highest area under the curve (AUC) and area under the precision-recall curve (AUPR) ratios across all distance groups in naive CD4 cells (Fig. 4a, b, Methods), whereas other methods performed comparably to random. Since SCENIC+ only provides final RE-TG pairs without relevance scores, we used the F1 score to assess its performance (Methods). scTFBridge outperformed SCENIC+ in F1 scores across all distance groups (Fig. 4c). In other cell types, due to the limited availability of eQTL pairs, all RE-TG pairs were treated as positive labels without further distance-based stratification. scTFBridge consistently outperformed baseline methods in both AUC and AUPR ratios across all evaluated cell types, achieving the highest overall ranking (Fig. 4d, e) and significantly higher F1 scores compared to SCENIC+ (Fig. 4f). Additionally, when using pcHi-C data as ground truth, scTFBridge also achieved the highest AUC across all distance groups in naive CD4 T cells (Fig. 4g).

**Fig. 4.**
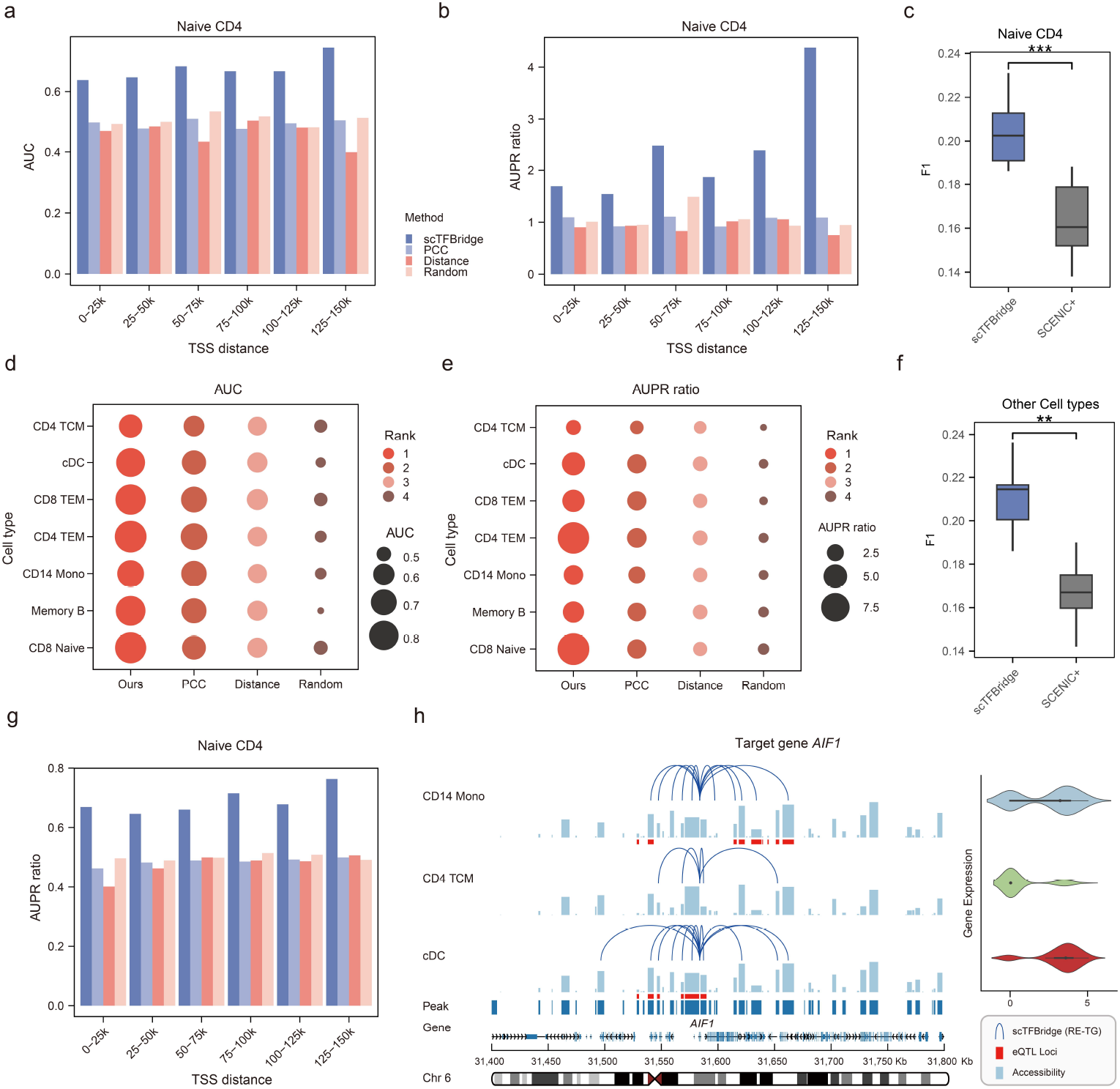
Benchmarking analysis of cell-type-specific cis-regulatory inference in PBMCs. a, b Benchmark performance (AUC and AUPR ratio) of different methods for cis-regulatory inference in naïve CD4+ T cells, evaluated using eQTL data as ground truth. c, F1 score comparison between scTFBridge and SCENIC+ for cis-regulatory inference in naïve CD4 + T cells. Statistical significance was assessed using a paired Student’s t-test, with *** indicating a p-value < 0.001. d, e Benchmark performance (AUC and AUPR ratio) of different methods of cis-regulatory inference across other cell types f, F1 score comparison between scTFBridge and SCENIC+ in the cis-regulatory inference across other cell types. Statistical significance was evaluated with a paired Student’s t-test, where ** indicates a p-value < 0.01. g, Benchmark performance (AUC) of different methods in the cis-regulatory inference of naïve CD4+ T cells, using the pcHi-C data as ground truth. h, scTFBridge-identified cis-regulatory interactions of *AIF1* in CD14 Mono, CD4+ T central memory (TCM) cells, and conventional dendritic cells (cDC), supported by eQTL evidence.

To further demonstrate its utility, we examined cis-regulatory interactions identified by scTFBridge that align with known gene regulatory mechanisms across various cell types. For instance, *AIF1* gene expression varies among cell types, being highly expressed in CD14 monocytes and cDCs, but with lower expression in T and B cells. To illustrate scTFBridge’s ability to capture cell-type-specific cis-regulatory interactions for *AIF1*, we analyzed a 200 kb region upstream and downstream region of its TSS, annotating cell-type-specific peak accessibility, eQTL SNP loci, and inferred RE-TG pairs predicted by scTFBridge (Fig. 4h). Using CD4 TCM cells as a reference, scTFBridge successfully identified the majority of known cis-regulatory elements in CD14 Mono and cDCs, supported by single-cell eQTL data. Additionally, scTFBridge uncovered potential novel regulatory elements that may be specific to these cell types (Fig. 4h). These results underscore the heterogeneity of cis-regulatory mechanisms across cell types and highlight scTFBridge’s potential to enhance and refine cell-type-specific GRN inference.

### Benchmarking analysis of cell-type-specific trans-regulatory inference

To evaluate the single-cell trans-regulatory inference capability of scTFbridge, we leveraged pre-identified TF-TG regulatory pairs from Chromatin Immunoprecipitation sequencing (ChIP-seq) data in the Cistrome database as the ground truth^61^. Specifically, we compiled regulatory data for 17 TFs and their corresponding putative target genes across four blood cell types (Supplementary Table 2). We benchmarked scTFBridge’s performance against alternative methods, including GENIE3^16^, PIDC^15^, PCC, and SCENIC+^38^, for trans-regulatory inference in these blood cells. As a case study, we analyzed the trans-regulatory scores of STAT1 across all single cells (Fig. 5a, Fig. S3-4) and examined its paired trans-regulatory TGs in CD14 monocytes. In this context, scTFBridge demonstrated superior performance, achieving an AUC score of 0.776 (0.531–0.583 for the others), and an AUPR ratio of 2.654 (1.166–1.388 for the others) (Fig. 5b, c). Furthermore, scTFBridge consistently outperformed all baseline methods across the benchmarking dataset, achieving higher AUC scores and AUPR ratios for all TFs tested (Fig. 5d, e). Unlike GENIE3 and PIDC, which use only gene expression data, scTFBridge combines scRNA-seq and scATAC-seq data, demonstrating the advantages of multi-omics integration for improved trans-regulatory inference. Furthermore, scTFBridge surpassed the multi-omics model SCENIC+, highlighting its robustness in analyzing complex regulatory networks.

**Fig. 5.**
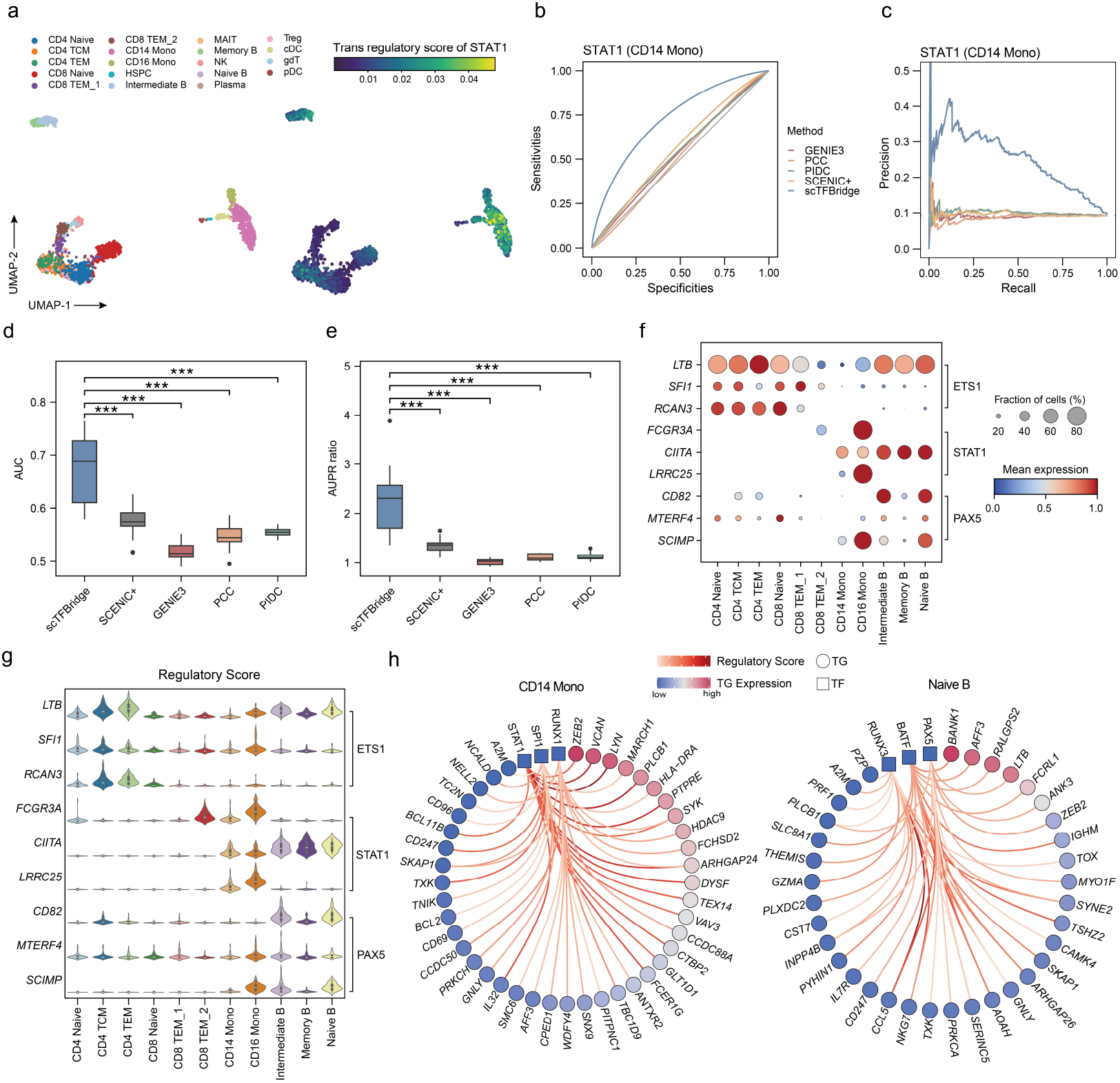
Benchmarking analysis of cell-type-specific trans-regulatory inference in PBMCs. a, UMAP visualization of modality-shared embedding of PBMCs. b, c Receiver operating characteristic curve and precision-recall curve of trans-regulatory potential inference of STAT1 in CD14 Mono cells. d, e Box plots comparing AUC and AUPR ratio scores across various methods for identifying putative TF target genes using ChIP-seq data. Statistical significance was evaluated with paired Student’s t-test, where *** indicates a p-value < 0.001. f, Top TF-regulated gene expression validated by ChIP-seq data across different cell types, including ETS1 in CD4+ T cells, STAT1 in CD14+ monocytes, and PAX5 in naïve B cells. g, Violin plots showing cell-level trans-regulatory scores of specific TF-TG pairs across different cell types. h, Network representations of inferred TF-TG pairs in CD14 Mono and naïve B cells. Node color corresponds to gene expression levels, while edge color represents regulatory scores.

Next, we explored trans-regulatory associations in other single-cell groups. Based on annotated TF-TG associations, we selected ETS1, STAT1, and PAX5 as the dominant regulatory TF in CD4 T cells, monocytes, and B cells, respectively, along with their most associated TGs. Comparing TG expression levels revealed cell-type-specific patterns of gene expression for these TF-TG pairs (Fig. 5f). We further evaluated the regulatory scores predicted by scTFBridge and found that the model effectively captured cell-type-specific trans-regulatory associations (Fig. 5g). To visualize these relationships, we reconstructed the inferred trans-regulatory networks for cell-type-dominant TFs and their associated TGs. For instance, we identified the STAT1*-ZEB2* regulatory link in CD14 Monocytes and BATF*-BANK1* regulatory link in B cells (Fig. 5h). These results underscore scTFBridge’s ability to resolve cell-type-specific trans-regulatory activities at single-cell resolution.

### scTFBridge uncovers cell-type-specific gene regulations in lung cancer susceptibility

To explore disease-specific single-cell gene regulation, we applied scTFBridge to a single-cell multi-omics dataset derived from lung cancer^62^. This dataset comprises paired scRNA-seq and scATAC-seq datasets collected from tumor-distant normal lung tissues of ever- and never-smokers (n = 8 per group). The modality-shared embedding of these single cells revealed a distinct separation based on annotated cell types from Long et al. (Fig. 6a). In contrast, the RNA-private and ATAC-private embeddings exhibited less cellular information, with the RNA-private embedding showing particularly reduced information content (Fig. S5).

**Fig. 6.**
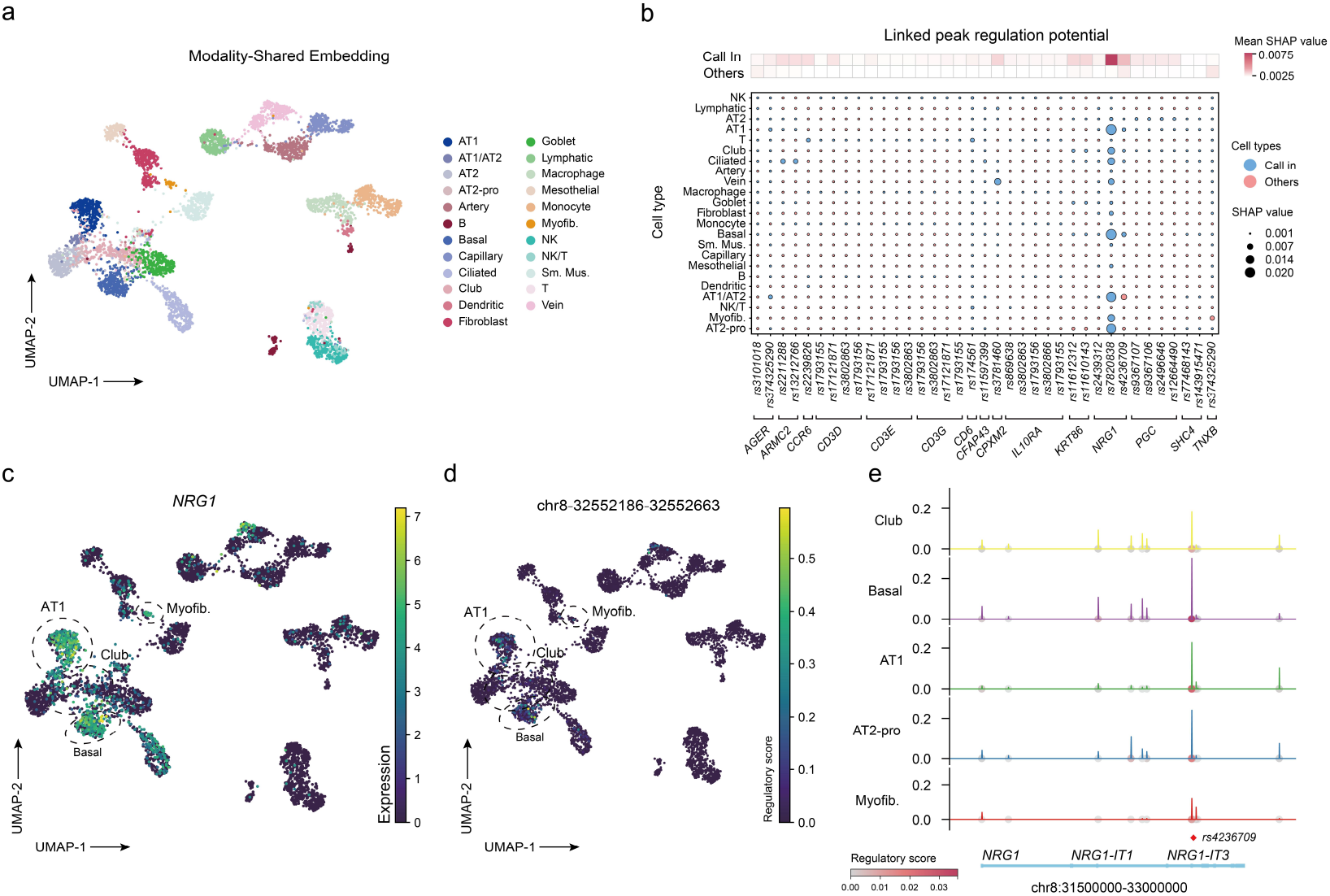
Cell-type-specific gene regulations in lung cancer susceptibility uncovered by scTFBridge. a. UMAP visualization of the modality-shared embedding of lung single-cell dataset. b. Regulation scores of tier-6 linkage variants and candidate cis-regulatory elements associated with target genes across different cell types. Dot color indicates cell-type categories, and dot size represents the regulation scores. c. UMAP visualization of *NRG1* gene expression of lung single-cell dataset as in (a). d. UMAP visualization of the regulation scores for the variant rs4236709-linked peak across different cell types as in (a). e. Visualization of peak accessibility and regulation scores within the *NRG1* locus across various cell types. Dot color corresponds to the regulatory score of a specific peak to *NRG1*.

Long et al. previously defined high-confidence RE-TG pairs (tier 6) that included lung cancer candidate causal variants (CCVs) colocalized REs and their TGs. This group proposed potential gene regulatory patterns contributing to lung cancer susceptibility. To validate scTFBridge’s ability to uncover cell-type-specific cancer susceptibility cis-regulations, we compared the inferred regulatory scores of identified RE-TG pairs across predefined “call in” cell subtypes and other cell types (Fig. 6b). The results demonstrated that scTFBridge effectively captures higher cell-specific cis-regulations for these RE-TG pairs.

Specifically, we visualized the expression of the target gene *NRG1* across all single cells and evaluated the regulatory score inferred by scTFBridge for a CCV-specific peak (chr8: 32552186–32552663) (fig. 6c). The findings confirmed that scTFBridge accurately reflects cell-type-specific cis-regulation. Furthermore, a comparison of the inferred regulatory scores across peaks within the *NRG1* gene region also indicated that the CCV-colocalized peak exhibited the highest regulatory potential (Fig. 6d, e). To further investigate context-dependent gene regulation, we extended our analysis to smoking-responsive CREs, the results confirmed reduced regulatory scores for selected RE-TG pairs in cells from the ever-smoker group (Fig. S6). These results underscore the capability of the scTFBridge model as a powerful tool for identifying gene regulatory events underlying human disease susceptibility.

## Discussion

Inferring dynamic gene regulatory processes at single-cell resolution advances our understanding of complex cellular regulation during development and disease, while also facilitating the discovery of genotype-phenotype associations^14,63,64^. Although multiple methods have been developed to infer GRNs using scRNA-seq or scATAC-seq datasets^38,65^, these approaches are often limited by the unpaired nature of the datasets and the high sparsity of single-cell data, making it challenging to accurately characterize complex cell-type-specific GRNs. The advent of single-cell multi-omics techniques allows for the simultaneous profiling of individual cells at both the transcriptomic (RNA-seq) and epigenomic (ATAC-seq) levels. This comprehensive characterization necessitates the development of novel computational methods tailored for single-cell multi-omics profiling. These datasets facilitate learning joint embeddings of single-cell features^2^, establishing associations between REs and TG expression^66^, and inferring GRNs^35,38^. Additionally, advances in biological deep learning for single-cell omics further provide more accurate representations of single-cell data, enhancing interpretability and offering deeper insights into cellular regulatory mechanisms^17,25,67^.

In this study, we introduced scTFBridge, an interpretable deep generative model designed to infer GRNs from single-cell multi-omics data. Leveraging mutual information theory, scTFBridge effectively separates the latent variable space into modality-shared and modality-private subspaces, addressing key challenges posed by modality gaps and information redundancy in multi-omics integration^44,68^. Furthermore, scTFBridge incorporates prior knowledge of TF-motif binding to regularize the biological learning of modality-shared embedding. This approach enables the model to capture TF-RE interactions within the shared embeddings, enhancing interpretability. Notably, each latent variable in the modality-shared embedding corresponds to a specific TF, forming a “bridge” between RNA-seq and ATAC-seq data to support accurate GRN inference. This architecture not only improves GRN inference accuracy but also ensures that the representations remain biologically interpretable.

We applied scTFBridge to single-cell multi-omics datasets derived from BMMCs^55^, rheumatoid arthritis cells^56^, PBMCs, and lung cells^62^ to evaluate its performance across diverse gene regulatory tasks. Leveraging a deep-learning model interpretation framework, scTFBridge effectively identified cell-type-specific TFs and uncovered the contributions of various TFs and REs to TG regulation in different cell types. Using promoter-capture Hi-C (pcHi-C), eQTL, and ChIP-seq data from blood as ground truth, scTFBridge demonstrated superior performance compared to baseline methods in inferring cis- and trans-regulatory interactions in the PBMC dataset. Furthermore, analysis of a lung cell multi-omics dataset demonstrated scTFBridge’s ability to identify candidate susceptibility genes associated with causal variants within cell-specific REs. These results highlight scTFBridge’s efficacy in disentangling complex regulatory features, thereby advancing single-cell GRN inference.

Despite its contributions, this study has several limitations. Our benchmarking analysis was primarily conducted on blood-derived datasets due to the limited availability of standardized ground truth data for other tissues and cell types, which restricts the generalizability of the method. Additionally, our analysis concentrated on a simplified model of TF and RE binding, which may limit its applicability in more complex TF regulatory contexts. While scTFBridge effectively extracts latent variables for RNA-seq and ATAC-seq data, it does not account for potential regulatory mechanisms involving microRNAs, non-coding RNAs, or private Peaks, which require further exploration and the development of complementary methodologies^2^.

In summary, scTFBridge provides a computational framework for inferring both cis- and trans-regulatory GRNs from single-cell multi-omics data. The core principle of biologically informed multi-modal disentanglement can be adapted to other single-cell multi-omics applications. This work emphasizes the importance of developing interpretable, biologically grounded multi-modal integration methods that advance our understanding of molecular regulatory processes at the single-cell level.

## Methods

### Framework of scTFBridge

scTFBridge is a deep learning-based computational framework designed to integrate single-cell multi-omics data and infer cell-type-specific GRNs. The framework utilizes a hybrid multimodal VAE architecture to decompose the data into shared and private latent spaces. The shared latent space captures regulatory interactions among TFs, REs, and TGs, and the private latent space encodes modality-specific information, such as non-TF regulatory factors or other cellular-specific features unique to each data modality.

RNA-seq and ATAC-seq data from the same cells are processed through separate encoders to generate shared and private latent distributions. A product of experts (PoE) module combines information from both modalities to form a joint shared distribution in the shared latent space. Both shared and private latent distributions are modeled as Gaussian distributions, with Kullback-Leibler (KL) divergence used for regularization. Latent representations sampled from these distributions are passed through modality-specific decoders to reconstruct the original data.

To disentangle shared and private latent spaces, scTFBridge incorporates a cross mutual information (CMI) block. This block minimizes mutual information between shared and private embeddings while maximizing mutual information between shared RNA-seq and ATAC-seq embeddings. To enhance biological relevance, prior knowledge of TF-motif binding scores is used to link shared ATAC-seq embeddings to REs in the ATAC-seq decoder. The shared latent variables, representing TF regulatory activity, enable the quantification of contributions from specific TFs and REs to observed regulatory interactions. By computing SHAP values for ATAC-seq REs and TF latent representations to TGs, scTFBridge enables the inference of both cis- and trans-regulatory networks, providing a detailed view of cell-type-specific GRNs.

#### Disentangled Multi-Modal VAE

For a given cell *n*, the observed multi-omics input data is denoted as *x* = {*x*_*r*_, *x*_*a*_}, where *x*_*r*_ represents RNA-seq data and *x*_*a*_ represent ATAC-seq data. we define a low-dimension latent variable *z* = {*z*_*r*_, *z*_*a*_} to represent the observed latent omics. The latent space *z* is divided into modality-shared and private representations, factorized as *z*_*r*_ = {*z*_*sr*_, *z*_*pr*_}, *z*_*a*_ = {*z*_*sa*_, *z*_*pa*_}. For a single modality (e.g., RNA-seq), the goal of the deep generative model is to maximize the likelihood of the observed data conditioned on the latent variables. This likelihood is expressed as Eq. (1):

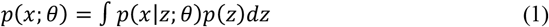

Here, *p*(*z*) is the prior distribution, typically set as a standard Gaussian distribution, and *p*(*x*∣*z*; *θ*) represents the process of generating the observed data from latent variable, and *θ* denotes the learnable parameters in the decoder. Direct computation of *p*(*x*;*θ*) is challenging, so we introduced a variational posterior *q*(*z*∣*x*;*ϕ*) to approximate the true poster distribution *p*(*z*∣*x*), where *ϕ* are the parameters of data encoder. Using this approximation, Eq. (1) can be rewritten as:

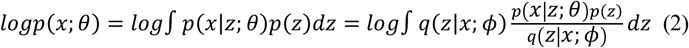

Using Jensen’s inequality, this leads to the evidence lower bound (ELBO):

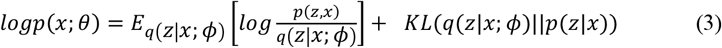

Since *KL*(*q*(*z*∣*x*; *ϕ*)∣∣*p*(*z*∣*x*)) ≥ 0, the ELBO is given by:

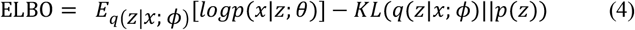

Here, *KL*(*q*∣∣*p*) represents the Kullback–Leibler divergence between *q*(*z*∣*x*;*ϕ*) and prior *p*(*z*). To maximize the log-likelihood, we can maximize the ELBO.

To disentangle the latent variables and address batch effects, the learning objective for each omics layer represents as follows:

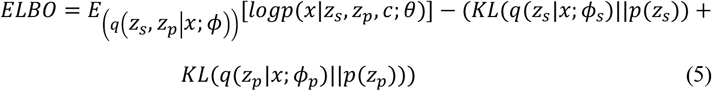

Here, *c* is the batch label used as a conditional variable in the generative process.

For the modality-shared representation, we use a product-of-experts (PoE) model to compute the joint variational posterior distribution for each omics modality:

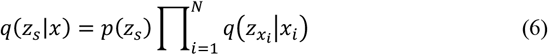

The prior *p*(*z*_*s*_) is modeled as a Gaussian distribution *N*(*z*_*s*_∣*µ*_0_, *δ*_0_), and each modality-specific posterior 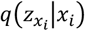 is also Gaussian, with mean 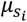 and variance 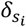. The joint distribution is defined by:

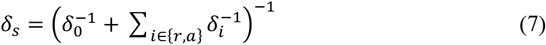

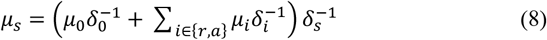

The overall learning objective for multi-modal integration during training is defined as:

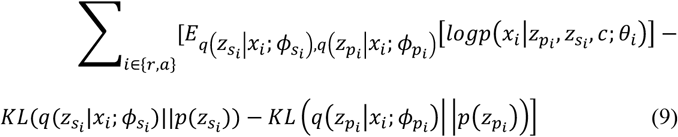

#### Cross mutual information Block

To achieve a well-disentangled multi-modal latent space, we incorporate mutual information (MI) theory to decouple shared and private representations. The loss function for this block is defined as in Eq. (10):

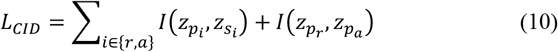

Here, 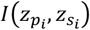 measures the MI between the private and shared latent embeddings within a single modality *i*, 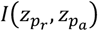 and measures the MI between private embeddings across modalities. The goal is to minimize the MI between modality-shared and private embeddings within a modality and between private embeddings across modalities. The MI between two variables *x* and *y* is defined as in Eq. (11):

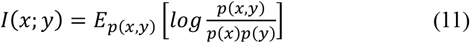

Since mutual information is generally intractable, we use an upper bound known as the Contrastive Log-ratio Upper Bound (CLUB). This approximation for *I*_*CLUB*_(*x, y*) is described as in Eq. (12):

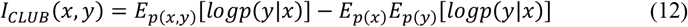

To validate that *I*_*CLUB*_(*x, y*) is an upper bound of *I*(*x*; *y*), we compute the difference Δ = *I*_*CLUB*_ − *I*(*x*; *y*) as in Eq. (13):

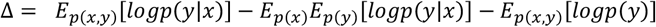

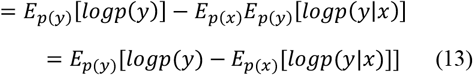

Using the marginal distribution *logp*(*y*) = *log*∫ *p*(*y*∣*x*)*p*(*x*)*dx* = *log*(*E*_*p*(*x*)_[*p*(*y*∣*x*)]) and Jensen’s inequality, we derive:

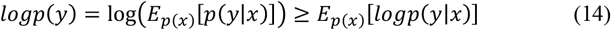

This shows that Δ ≥ 0, confirming that *I*_*CLUB*_(*x, y*) is indeed an upper bound of *I*(*x*; *y*).

Given a dataset of *N* sample pairs 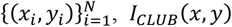 can be computed as:

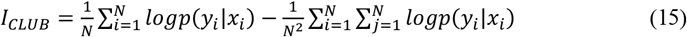

This simplifies to:

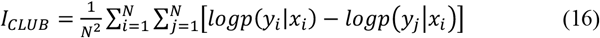

We employ a nonlinear neural network to estimate *p*(*y*∣*x*), with the log-likelihood computed as:

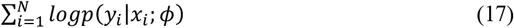

During each training epoch, the parameters of the distribution estimator are first updated to maximize Eq. (17), improving the accuracy of conditional distribution estimation. Subsequently, other parameters of scTFBridge are updated to minimize *I*_*CLUB*_, thereby reducing the mutual information between the two variables.

#### TF-Binding informed ATAC Decoder

To ensure that the joint shared embedding represents transcription factor (TF) regulatory activities, we incorporated prior knowledge of TF-motif binding scores into the architecture of the ATAC-seq decoder. We utilized 713 TF position weight matrices (PWMs) for known motifs, sourced from the PECA2 GitHub repository (https://github.com/SUwonglab/PECA). This repository aggregates motif data from widely used databases, including JASPAR^69^, TRANSFAC^70^, and UniPROBE^71^. From the single-cell RNA-seq dataset, the 128 most highly expressed TFs were selected as dimensions of the modality-shared embedding. Each TF corresponds to a single latent variable in the shared embedding. Regulatory elements (REs) were scanned for TF binding motifs using HOMER. This produced a binding score matrix *B*, which quantifies the interaction strength between each TF and the REs.

The ATAC reconstruction decoder is implemented as a single-layer neural network. The decoder receives the combined latent variables as input and reconstructs ATAC-seq signals (peaks) constrained by biological priors. The binding score matrix *B* serves as a mask to constrain the weight matrix *W* of the decoder. *B* ensures that connections from TF latent embeddings to ATAC-seq peaks reflect known TF-RE interactions. For each connection in the decoder, the weight matrix *W* is modulated by the binding score matrix *B*, as follows:

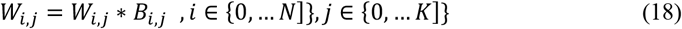

Where *N* is the number of TF dimensions and *K* is the number of REs in ATAC-seq.

#### Cross modal inference

In addition to modality-private self-reconstruction, the scTFBridge training process incorporates cross-modal generation to learn cis-regulatory relationships between ATAC-seq and RNA-seq. This feature enables inference across modalities, allowing the model to predict missing data in one modality using information from the other. When RNA-seq data is missing, the Product-of-Experts (PoE) approach is employed to use the shared latent distribution of ATAC-seq 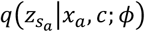 as the multi-modal joint distribution for cross-omics generation. The missing modality-private latent variable 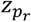 is reparametrized from a prior normal Gaussian distribution 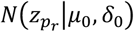. Using 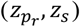, the model recovers the RNA-seq data by optimizing the following objective:

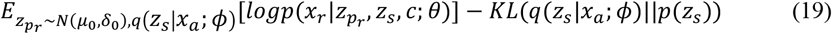

Similarly, we can derive the objective function for generating ATAC-seq data from RNA-seq:

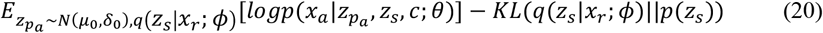

#### Learning Object

In general, the final optimizing objective of scTFBridge can be expressed as follows:

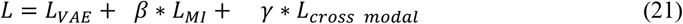

The first term optimizes the disentangled VAE for learning omics-specific representations and accurate data reconstruction, it measures how well the model reconstructs each modality from the latent representations and regularizes the latent space by minimizing the divergence between the latent distributions and the prior Gaussian distribution. The second term evaluates and promotes the disentanglement of the shared and private latent representations by minimizing the MI between them. The third term ensures that the model can perform effective cross-modal inference, using the shared and reparametrized latent variables. These terms collectively ensure that scTFBridge achieves biologically meaningful integration, disentanglement, and cross-omics prediction. Hyperparameters are used to balance the weights of each learning objective, with example values for each dataset provided in Supplementary Table 3.

### Single cell multi-omics data preprocessing

In this study, we implemented a single-cell multi-omics data preprocessing workflow using the Scanpy package^72^. For the scRNA-seq data, TFs were isolated, normalized, and log-transformed. We then selected the top 128 highly variable TFs for focused embedding in the shared TF space. Additionally, the top 3,000 highly expressed genes were identified as (TGs). These TGs were normalized and log-transformed to serve as input for scTFBridge. For scATAC-seq data, chromatin accessibility peaks were binarized and filtered to retain peaks with open accessibility in at least 1% of all cells. To ensure accurate integration, only cells represent in both datasets were included in the analysis.

### SHAP value for cell-level and cell-type-specific GRN inference

SHAP values provide a robust framework for quantifying the contributions of features in deep learning models and are widely applied in biologically interpretable biomarker discovery. Building on prior studies^54^, we utilized reconstruction loss between original and imputed expression values of TGs as the objective to compute SHAP values for input features, focusing on their contribution to the reconstruction rather than direct expression effects. The strength of each feature’s contribution to TGs was quantified as the average of the absolute SHAP values across cell samples in specific cell groups.

For cis-regulatory contributions, we calculated regulatory elements (REs) associated with TGs via cross-omics generation. The cis-regulatory score for a RE-TG pair was defined as:

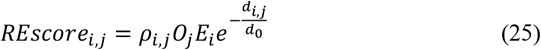

Where *ρ*_*i,j*_ is the SHAP value of RE *j* and TG *i*; *O*_*j*_ is the average chromatin accessibility of this cell type; *E*_*i*_ is the average expression of TG *i* in the cell type; *d*_*i,j*_ is the distance between genomic locations of *j* and *i* ; and *d*_0_ is the normalization factor for genomic distance, set to 30,000 bp in this study.

For trans-regulatory contributions, we quantified the influence of each TF variable in the shared embedding space on TGs. The trans-regulatory score was directly defined as the SHAP value:

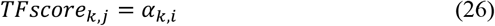

Where *α*_*k,i*_ represent the SHAP value quantifying the contribution of TF *k* to TG *i*.

To infer regulatory networks at the individual cell level, we extended the computation of SHAP values to account for unique cell-level characteristics. Instead of averaging values across a specific cell type, we calculated SHAP values using the average gene expression and chromatin accessibility across all cell types. This method enabled the reconstruction of cell-specific GRNs, providing a fine-grained view of regulatory interactions that account for each cell’s unique features.

### Latent Embedding Contributions Estimation

To evaluate the contributions of modality-private and shared components in the generative process of RNA-seq data from latent variables, we employed SHAP values. For each latent dimension, the average of the absolute SHAP values was computed across all TGs to derive global contribution scores. The overall contributions of the private and shared embedding spaces were then estimated by summing the contribution scores of all latent variables within their respective spaces. To ensure comparability, the two contribution scores (private and shared) were normalized to the range [0,1] by dividing each score by the combined sum of both scores. These normalized proportions were then ranked to determine the relative influence of private and shared components on gene regulation.

Using these normalized contribution proportions, we identify genes primarily regulated by private information as opposed to shared features and highlighted genes more influenced by shared features relative to private components. To further explore the biological relevance of these findings, we selected the top 100 ranked genes from each group for Gene Ontology (GO) enrichment analysis. Using the gprofiler2^73^ website, we examined enriched categories across biological processes pathways, gaining insights into the distinct regulatory roles of private and shared latent spaces in cellular functions.

### Benchmarking analysis of GRN inference

To evaluate the performance of scTFBridge in predicting cis- and trans-regulation, we conducted a benchmarking analysis against established methods using consistent dataset splits to ensure fairness in comparison. For cis-regulation prediction, we compared scTFBridge with PCC, TSS distance, random prediction, and SCENIC+^38^. For trans-regulation prediction, we selected PCC, SCENIC+^38^, GENIE3^16^, and PIDC^15^ as baseline methods.

### Heritability enrichment analysis

We utilized stratified linkage disequilibrium score regression (S-LDSC)^57^ to assess the heritability enrichment of various peak sets. First, we converted the genomic coordinates of peaks from hg38 to hg19 to align with the GWAS summary statistics of RA. SNPs with a minor allele frequency (MAF) ≥ 0.05 associated with the peak sets of interest were selected. Each 200 bp peak window was extended by 500 bp on either side, resulting in a 1.2 kb genomic interval per peak. Using S-LDSC, we then calculated population-specific linkage disequilibrium (LD) scores and quantified the proportion of RA heritability attributable to modality-shared and modality-private peak sets from different single-cell populations in the RA dataset, as analyzed using scTFBridge.

### Evaluation metric

#### AUPR Ratio

The AUPR Ratio measures the improvement of a predictor over random chance in terms of precision-recall performance. It is defined as the ratio of the Area Under the Precision-Recall (AUPR) curve of a method to that of a random predictor. For a random predictor, the AUPR is equal to the fraction of positive samples in the dataset *P*, which is defined as:

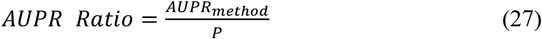

#### AUC

The AUC score evaluates the performance of a predictor in distinguishing between positive and negative samples. It is derived from the Receiver Operating Characteristic (ROC) curve, which plots the true positive rate (sensitivity) against the false positive rate (1-specificity) across thresholds.

#### F1 Score

The F1 Score assesses the balance between precision and recall, particularly useful when dealing with imbalanced datasets. It is computed as the harmonic mean of precision and recall, which is defined as:

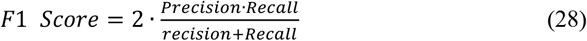

### Implementation Details of scTFBridge

We implemented scTFBridge using the PyTorch deep learning library in Python. The model was trained with the Adam optimizer, employing a learning rate of 1e-4 and weight decay of 1e-6, both reduced by epoch. Model training was conducted over 150 epochs with early stopping triggered after 10 steps of no improvement. To ensure robustness, the datasets were divided into five folds for cross-validation, with each fold split into 64% for training and 16% for testing. The testing set was used to identify the best model checkpoint and optimize hyperparameters, while SHAP values for GRN inference were computed using the validation datasets. All experiments were conducted on a workstation equipped with two Intel Xeon Silver 4210R CPUs and eight NVIDIA GeForce RTX 4090 GPUs.

## Data Availability

All datasets used in this study are publicly accessible. The BMMC single-cell multi-omics dataset was downloaded from the GEO database with access code GSE194122. The PBMC single-cell multi-omics data was obtained from the 10x Genomics (https://support.10xgenomics.com/single-cell-multiome-atac-gex/datasets/1.0.0/pbmc_granulocyte_sorted_10k) and the preprocessed data was download from the website(https://scglue.readthedocs.io/zh-cn/latest/data.html). The rheumatoid arthritis multi-omics data is available on the GEO database under access code GSE243917. The lung single-cell multi-omics data is deposited in the GEO database under access code GSE2414468. eQTL profiling data can be downloaded from the scQTLbase^74^ website (http://bioinfo.szbl.ac.cn/scQTLbase/Home/), with the specific file IDs of sc-eQTLs used in this study detailed in Supplementary Table 1. TF regulatory scores from ChIP-seq data were collected from Cistrome (http://cistrome.org/db), with the associated Cistrome IDs provided in Supplementary Table 2. The GWAS statistic profile of rheumatoid arthritis can be found at https://www.ebi.ac.uk/gwas/studies with accession number GCST90132222.

## Code availability

The scTFBridge model was constructed using Python, while R and Python were employed for data visualization. S-LDSC software(https://github.com/bulik/ldsc) was used to process GWAS summary statistics and disease heritability enrichment. The deep learning interpretation module was implemented on the source code of the previous study (https://github.com/suinleelab/PAUSE). All source code utilized in this study is publicly available on GitHub (https://github.com/FengAoWang/scTFBridge).

## Acknowledgments

This study was supported by the National Key R&D Program (2023YFF1204701, 2022YFF1202101), the Self-supporting Program of Guangzhou Laboratory (SRPG22007), the CAS Research Fund (XDB38050200), Guangdong Basic and Applied Basic Research Foundation (2023B1515130008). We thank technical support from the Data Science Platform of Guangzhou National Laboratory and the Bio-medical Big Data Operating System (Bio-OS).

## Author contributions

Y.L., J.L., and R.H. conceived the project. J.L. and A.W. designed the framework and loss designs. J.L. and A.W. wrote the manuscript with contributions from all authors. Y.L. supervised the entire project. All authors read and approved the final manuscript.

